## Supplemental Figures for "The Rhizobial effector NopT targets Nod factor receptors to regulate symbiosis in *Lotus japonicus*"

**Supplementary Files**

**Supplemental Figures**

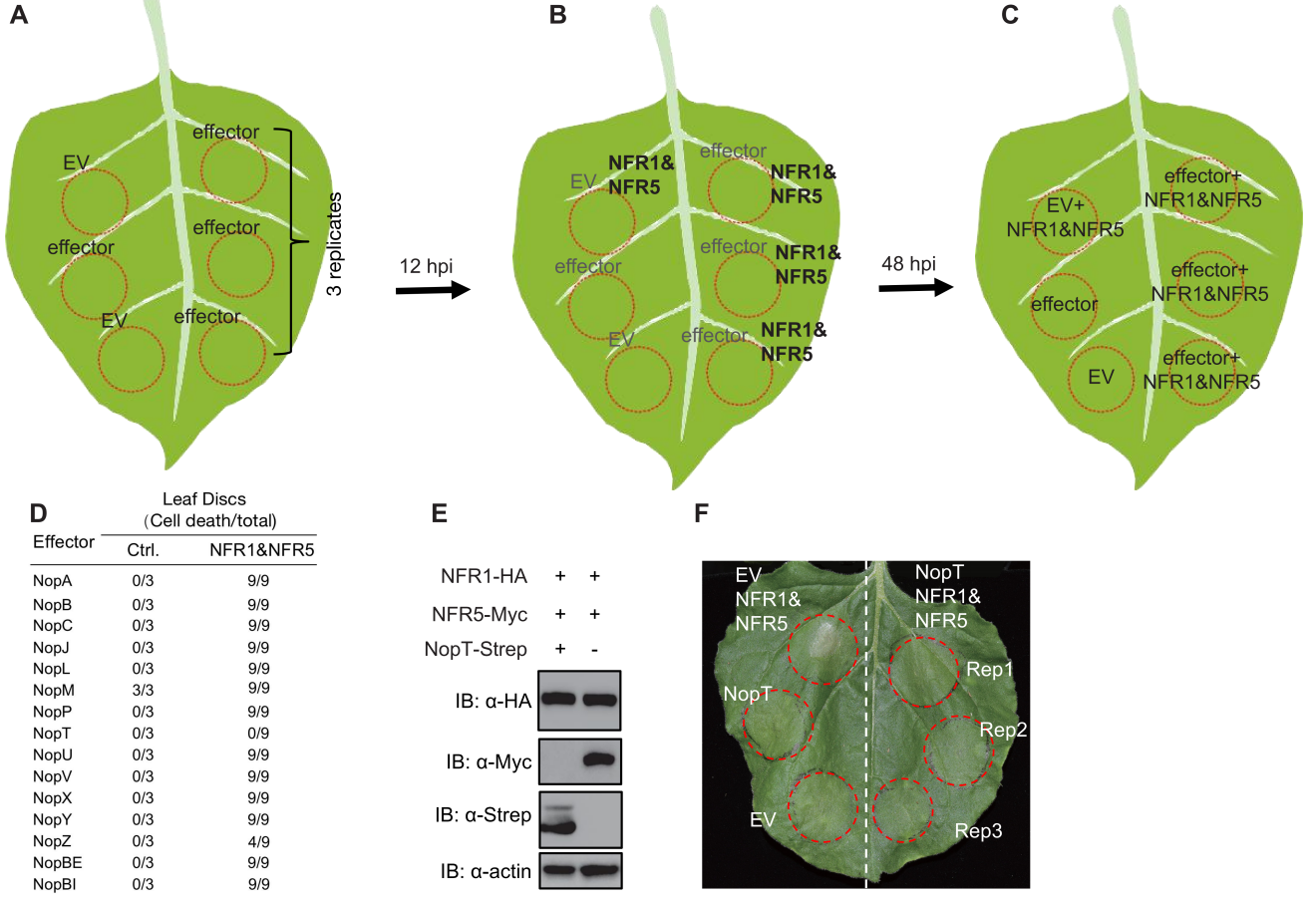

Supplemental Fig. 1. Analyses of the functions of all 15 effectors from *S. fredii* NGR234 in preventing NFR1- and NFR5-induced cell death in *N. benthamiana* leaves.

15 genes from *S. fredii* NGR234 encoding effector proteins were amplified and cloned into binary vector for expression of Strep-tagged proteins in *N. benthamiana*. NFR1 and NFR5 from *L. japonicus* were C-terminally tagged with HA and Myc, respectively, and cloned into binary vectors for expression in *N. benthamiana*. All plasmids were electroporated into *Agrobacterium tumefaciens* EHA105.

(A) *Agrobacterium* strain harboring each effector gene or control empty vector (EV) were infiltrated into *N. benthamiana*. 12-hour post infiltration (hpi), *agrobacterium* strains harboring NFR1 and NFR5 were mixed and hand-infiltrated in the same leaf discs as shown in (B). 48 hpi, cell death phenotypes were evaluated. (C) At least three replicates in the same leaf were performed to test the function of each effector in suppressing cell death-triggered by NFR1 and NFR5. (D) Results of all 15 effector proteins in suppressing programmed cell death by overexpression of NFR1 and NFR5. (E) Abundance of NFR1, NFR5, and NopT proteins, as measured by immunoblot with specific antibodies. Leaf discs expressing NFR1, NFR5, and NopT or NFR1 and NFR5 and with EV were used for immunoblot analyses. Actin was used as loading control. (F) Another example of suppression of programmed cell death by NopT in the leaf discs co-expressing with NFR1 and NFR5. Rep1, Rep2, and Rep3 represent three technical repeats in the same leave.

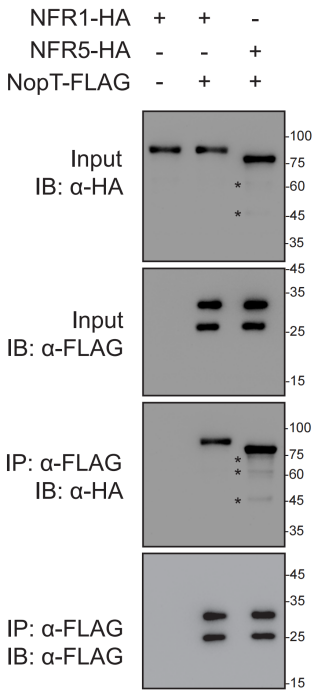

**Supplemental Fig. 2. NopT interacts with both NFR1 and NFR5.**

The full blot of co-IP assay as shown in Fig. 2D. The smaller bands are marked with asterisks. Compared to NFR1, NFR5 might be more susceptible to degradation during protein extraction from *N. benthamiana* leaves.

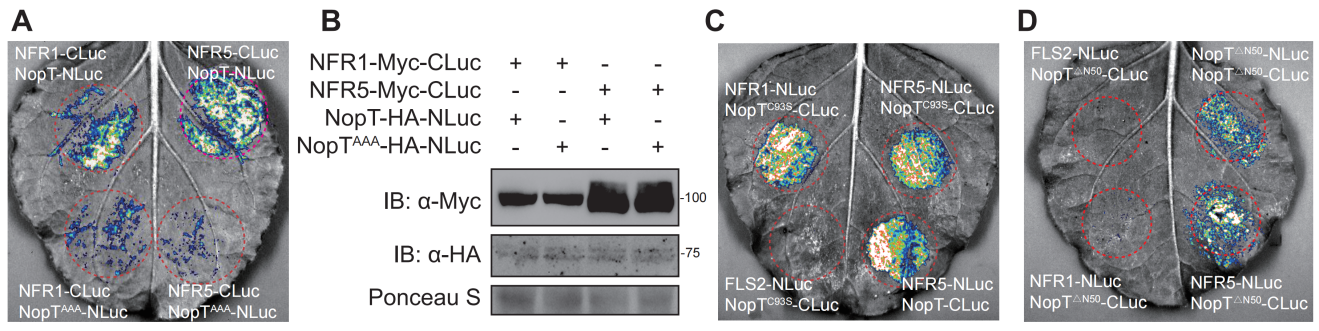

Supplemental Fig. 3. Split–LUC assay testing the interactions between different NopT mutants and NFRs.

(A) The interaction between NopT with mutated acylation site and NFR1/NFR5. NopT^G50A/C51A/C52A^ represents a modified NopT form lacking acylation sites. (B) The protein expressing of indicating genes in the leaves of (A). (C) Nop^C93S^ interact with NFR1 and NFR5. (D) The interaction between NopT^ΔN50^ and NFR5. NopT^ΔN50^ represents a truncated NopT with deletion of 50 amino acid residues from its N–terminus.

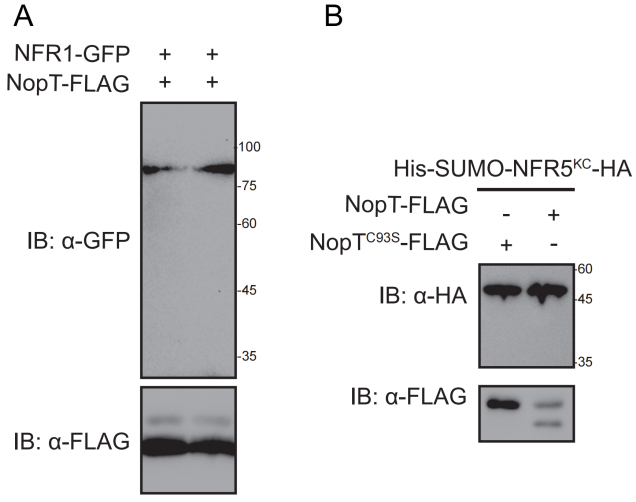

Supplemental Fig. 4. NopT cleaves NFR5 but not NFR1.

Proteins with indicated tags were expressed in either *N. benthamiana* or *E. coli* cells and detected by immunoblotting. (A) NopT did not cleave NFR1-GFP expressed in *N. benthamiana* leaves. (B) NopT expressed in *E. coli* cells did not cleave co-expressed SUMO-NFR5^KC^-HA. KC: Kinase domain and C-terminal domain, a modified NFR5^CD^ without the JM.

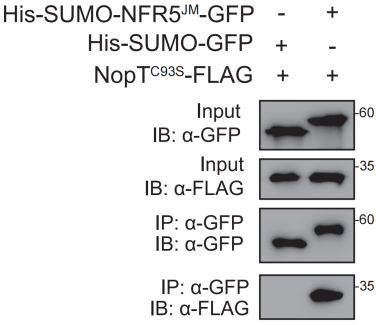

Supplemental Fig. 5. NopT interacts with the JM domain of NFR5.

NopT interacts with NFR5^JM^ in *in vitro* pull-down assay. IB: immunoblotting, IP: immunoprecipitation. His-SUMO-NFR5^JM^-GFP: a recombinant protein containing His tag, SUMO tag, the JM domain of NFR5, and GFP.

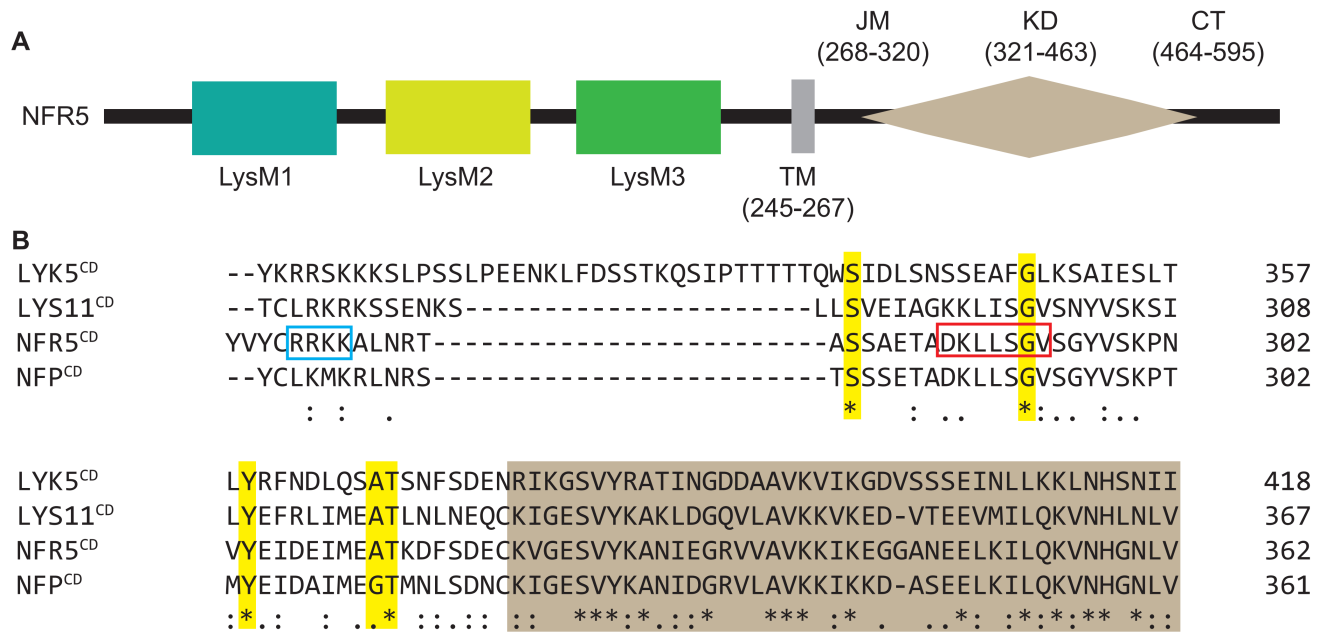

Supplemental Fig. 6. Conserved domains and residues of NFR5 and related proteins.

(A) Domain and motif annotation of NFR5. LysM: Lysin-Motif; TM: transmembrane domain; JM: juxtamembrane domain; KD: kinase domain; CT: C-terminal domain. (B) Alignment of amino acid residues of AtLYK5, LjLYS11, NFR5 and MtNFP close to the RRKK motif in the JM domain of NFR5. Five highly conserved residues marked by yellow coloration (S283, G294, Y304, A310, T311 in NFR5) and the brown box indicates the start region of kinase domain. The blue frame delineates the RRKK motif in NFR5 and the red-framed residues are similar to the NopT autocleavage region.

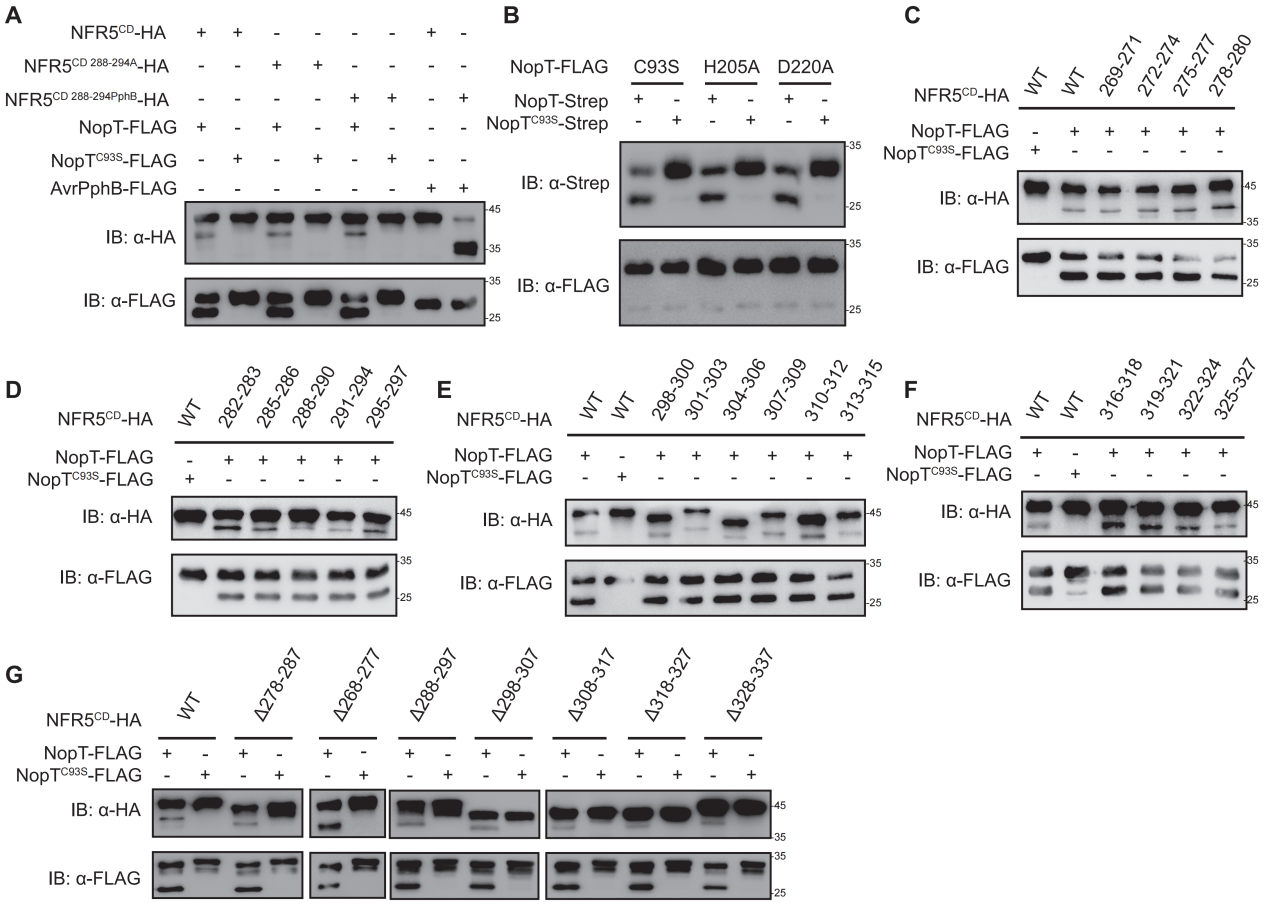

Supplemental Fig. 7. NopT cleaves NFR5 at the juxtamembrane domain.

(A) Cleavage assay using two mutant versions of NFR5^CD^ in the presence of NopT or AvrPphB. NFR5^CD 288-294A^ represents a mutant NFR5^CD^ with residues 288 to 294 replaced with seven alanines; NFR5^CD 288-294PphB^ represents a mutant NFR5^CD^ with residues 288 to 294 replaced with AvrPphB recognition sites. (B) Autocleavage assay, using different NopT variants with single amino acid mutations. H205A and D220A indicate His-205 and Asp-220 replaced with alanine, respectively. (C-F) NopT cleavage assay using 19 variants of NFR5^CD^ with three adjacent amino acids replaced with three alanines. The numbers indicate the location of the three residues replaced with alanines. (G) NopT cleavage assay using seven truncated variants of NFR5^CD^ with 10 adjacent amino acids deleted. The numbers indicate the location of the 10 deleted residues.

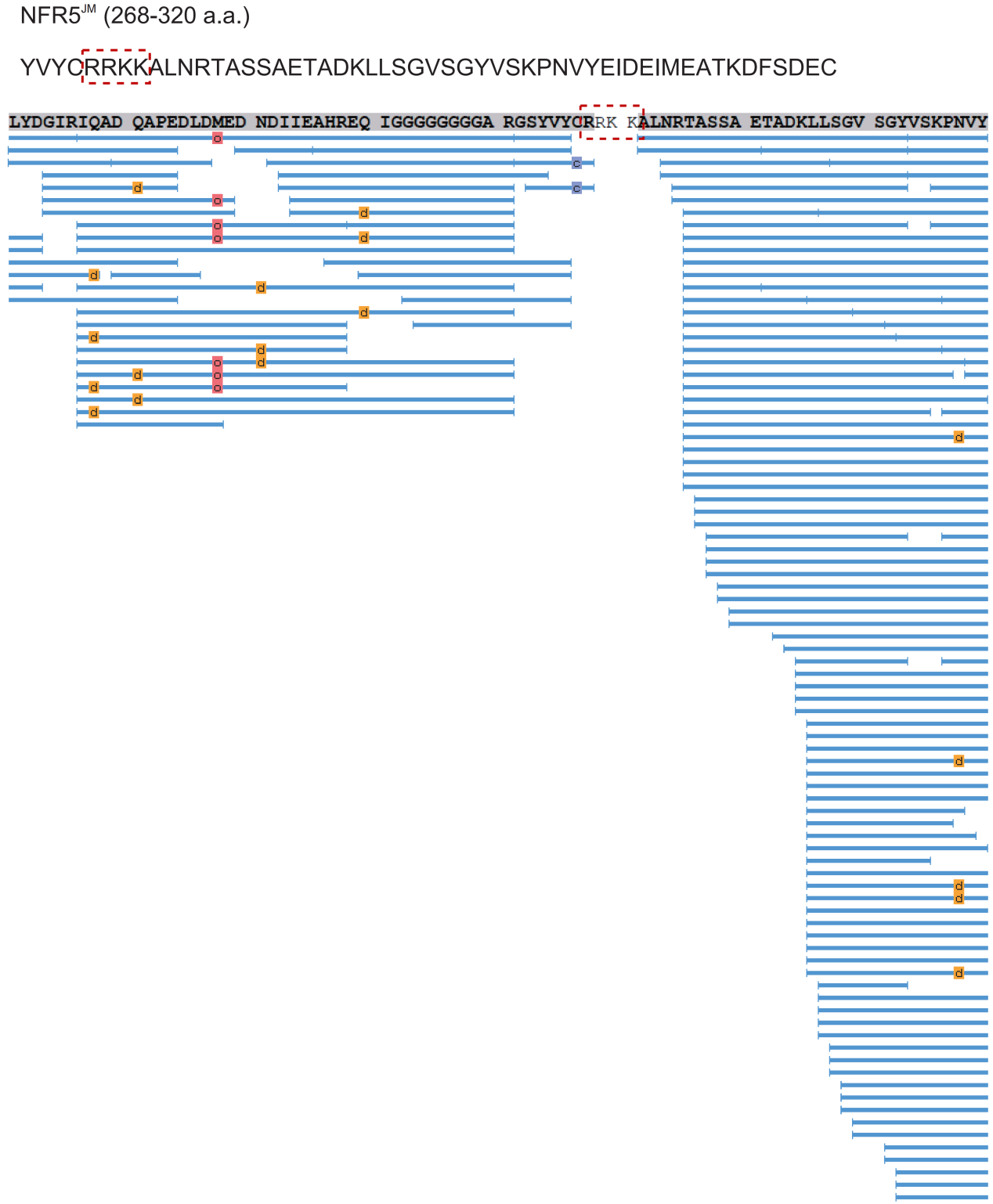

Supplemental Fig. 8. Mass spectrometry analysis of cleavage site of NFR5 by NopT.

(A) The sequence of NFR5 juxtamembrane domain (286-320 a.a.). (B) Mass spectrometry analysis of peptides of NFR5^CD^ after proteolysis by NopT. Blue lines indicated the peptides characterized by Mass spectrometry. Red frame delineated cleavage site of NFR5 identified using Mass spectrometry.

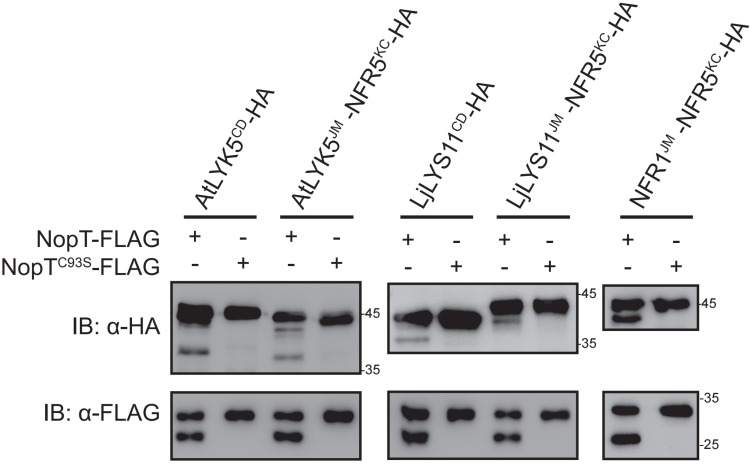

Supplemental Fig. 9. NopT cleaves the NFR5 homolog proteins at the juxtamembrane domain.

NopT but not NopT^C93S^ could cleave the CDs of AtLYK5 and LjLYS11 expressed in *E. coli* cells. NopT could cleave both AtLYK5^JM^–NFR5^KC^, LjLYS11^JM^–NFR5^KC^ and NFR1^JM^–NFR5^KC^, three recombinant proteins expressed in *E. coli* cells. AtLYK5^JM^–NFR5^KC^, LjLYS11^JM^–NFR5^KC^, and NFR1^JM^–NFR5^KC^ recombinant NFR5 proteins with the the JM was replaced with the JMs from AtLYK5, LjLYS11 and NFR1, respectively. KC: Kinase domain and C-terminal domain as ilustrated in Fig. S4B.

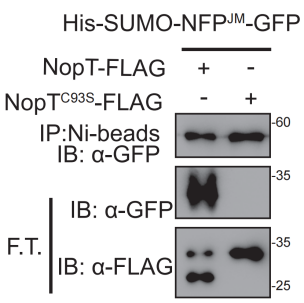

Supplemental Fig. 10. NopT cleaves the JM of MtNFP.

NopT cleaves a recombinant protein where His–tagged SUMO and GFP is bridged with the JM domain of NFP. F.T. represents flow through sample after Ni–beads purification. IB: immunoblotting.

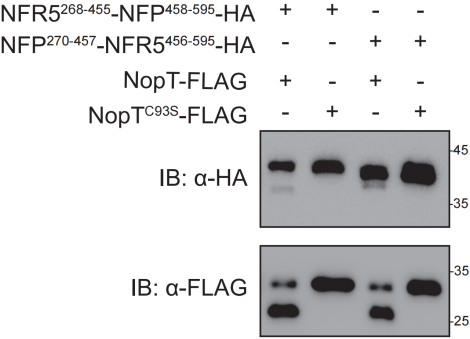

Supplemental Fig. 11. NopT cleaves recombinant proteins NFR5^268-445^-NFP^458-595^ and NFP^270-457^-NFR^5456-595^.

NopT but not NopT^C93S^ could cleave recombinant proteins NFR5^268-445^-NFP^458-595^ and NFP^270-457^-NFR^5456-595^ detected by immunobltting (IB).

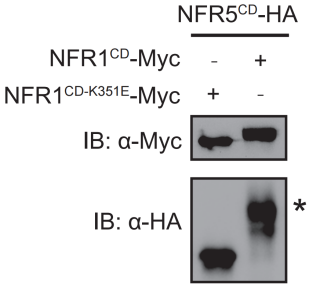

Supplemental Fig. 12. NFR1^CD^ phosphorylates NFR5^CD^ *in vitro*.

The cytoplasmic domains (CDs) of NFR1, NFR5, and kinase–dead NFR1^CD-K351E^ were used for kinase assays in *E. coli* cells. Proteins were detected by immunoblotting using anti–HA and anti–Myc antibodies. Asterisk indicated the band retardation on the gel representing the phosphorylated NFR5^CD^ proteins.

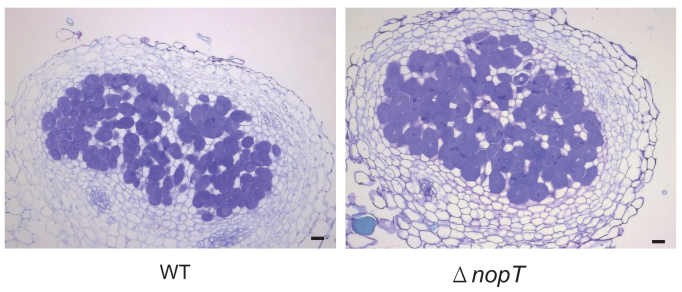

**Supplemental Fig. S13. Cross-section images of nodules at 14 dpi after toluidine blue staining.**

NGR234 and NGR234*ΔnopT* mutants inoculated *L. japonicus* Gifu at 14 dpi. Scale bars correspond to 100 μm.

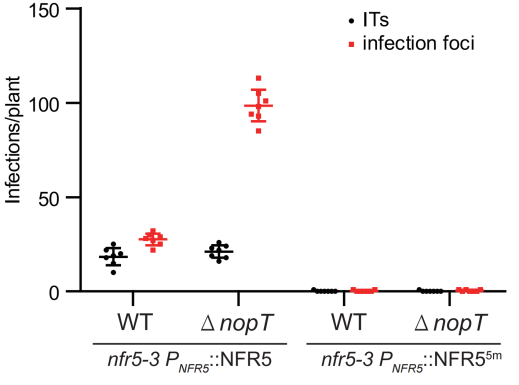

Supplemental Fig. 14. Rhizobial infection in the mutant versions of NFR5.

NFR5^5m^ failed to form rhizobial infection (n=7, Student’s *t*–test: P < 0.01). The transgenic roots expressing NFR5 or NFR5^5m^ in the *nfr5-3* mutant under the control of native promoter were inoculated with GFP-labelled wild type NGR234 and NGR234*ΔnopT* respectively. The data was collected 9 days post inoculation. NFR5^5m^ represents a mutant version of NFR5 with S283Y, G294Q, Y304S, A310I, T311Y point mutation.

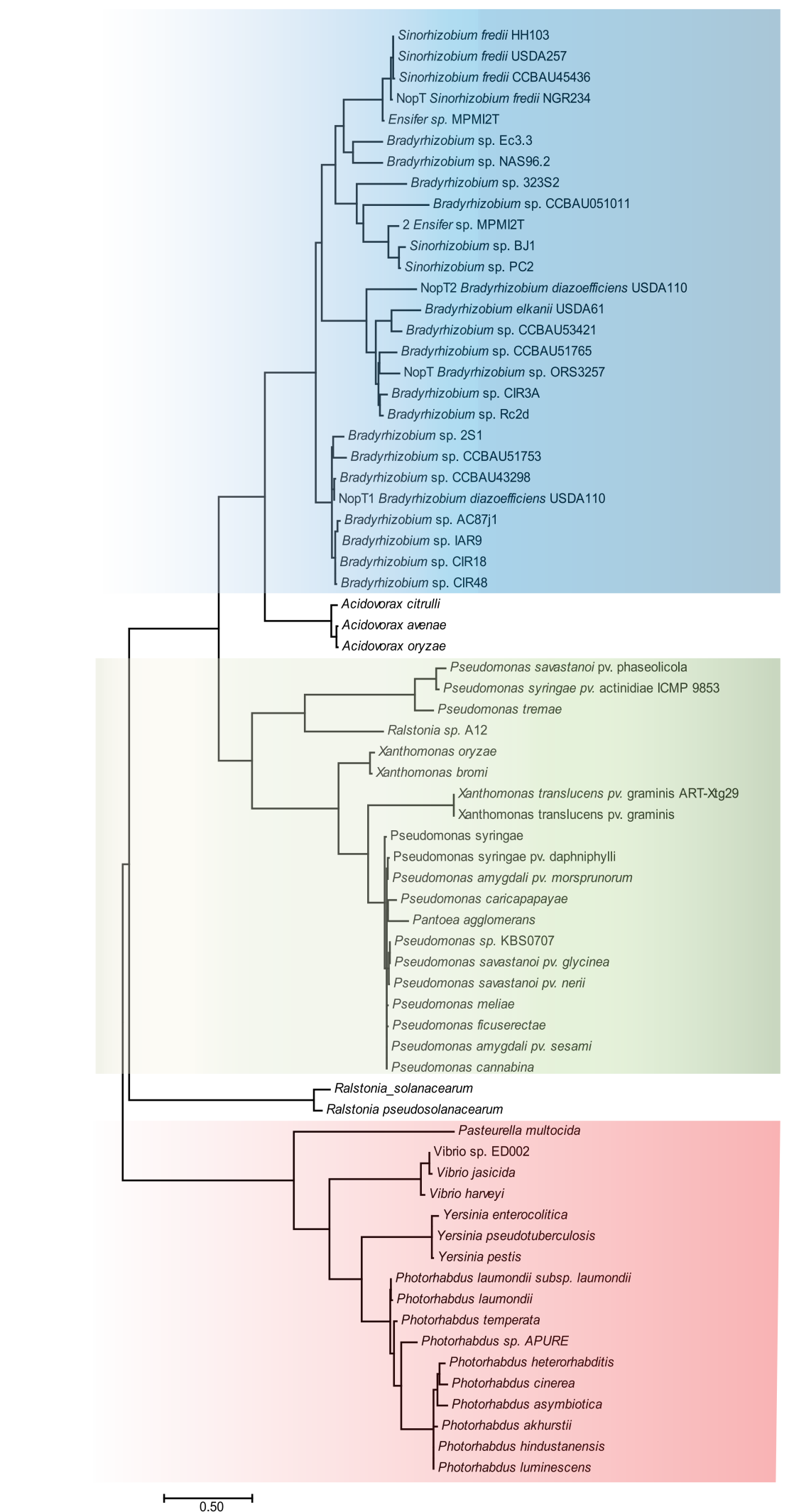

Supplemental Fig. 15. Phylogenetic tree based on the amino acid sequence of NopT homologs from different bacterial species.

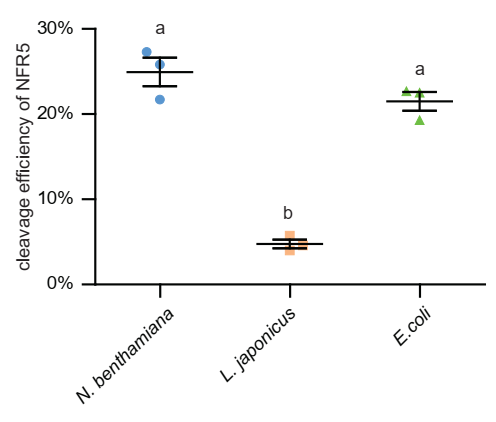

Supplemental Fig. 16. The cleavage efficiency of NFR5 in different system.

The relative cleavage efficiency for experiment displayed in Fig. 3A (*N. benthamiana*), Fig. 3B (*L. japonicus*) and Fig. 3D (*E. coli*) quantified from three biological replicates using ImageJ software. Values are means ± SE (n=3, Student *t*-test, letters represent significant differences, p<0.05).

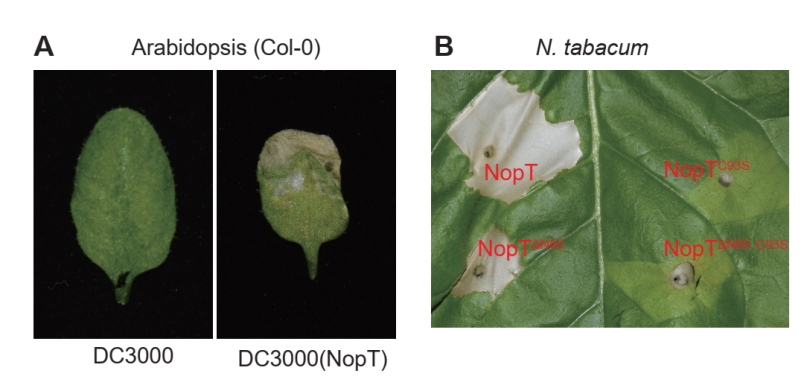

Supplemental Fig. 17. Transient expression of NopT triggers cell death in *Arabidopsis thaliana* and *Nicotiana tabacum*.

(A) *Pseudomonas syringae* pv tomato DC3000 harboring control plasmids (left) or expressing NopT (right) were infiltrated into *Arabidopsis* leaves. Pictures were taken three days post inoculation. (B) *Agrobacterium* strains harboring *NopT*, *NopT* with an N-terminal 50–amino acid deletion, *NopT^C93S^*, or *NopT^C93S^* with an N-terminal 50–amino acid deletion were infiltrated into *N. tabacum* leaves. Pictures were taken three days post inoculation.

**Supplemental Table 1. Phosphopeptides identified by liquid chromatography–mass spectrometry.** An *in vitro* kinase assay was performed using the CD of NFR1 and NopT. NopT was separated by SDS-PAGE, and the digested gel slices representing the phosphorylated NopT were used for analysis. The deduced phosphopeptides are listed.

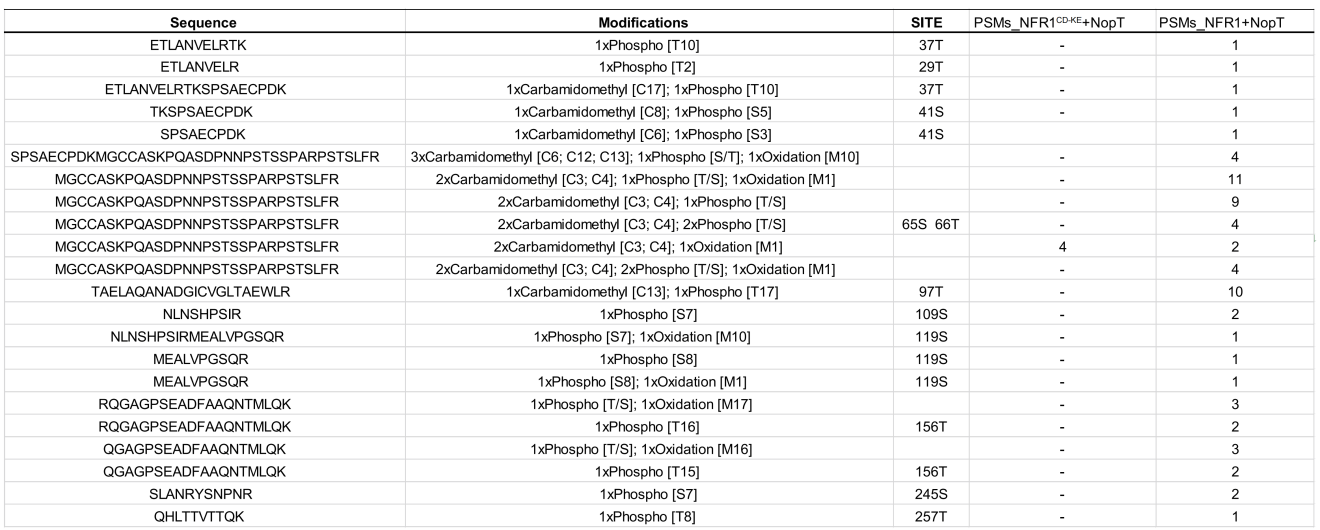

**Supplemental Table 2. Primers used in this study.**

| **Used for** | **Target genes** | **Names** | **Sequences (from 5’ to 3’)** |
| --- | --- | --- | --- |
| Overexpression in Nicotiana benthamiana | NopT or NopT^C93S^ | p5X-NopT-F | agaacacgggggactctagaatgcacagtcccatcag |
|  |  | p5X-NopT-R | ccgctgttatcggtacctgtcatcttttgggtggtcac |
|  | NopT^ΔN50^ | p5X-NopTdeN50-F | agaacacgggggactctagaatgtgctgcgccagcaag |
|  | NFR1 | p5X-NFR1-F | agaacacgggggactctagaatgaagctaaaaactggtctact |
|  |  | p5X-NFR1-R | ccgctgttatcggtacctcttctcacagac |
|  | NFR5 | p5X-NFR5-F | agaacacgggggactctagaatggctgtcttctttcttacc |
|  |  | p5X-NFR5-R | ccgctgttatcggtaccacgtgcagtaat |
|  |  | p5X-NFR5gfp-R | tcccgggagcggtaccacgtgcagtaatggaagt |
|  | AvrPphB | p5X-AvrPphB-F | agaacacgggggactctagaatgaaaataggtacgcaggcc |
|  |  | p5X-AvrPphB-R | ccgctgttatcggtacccgaaactctaaactcgtttacgc |
| templates for protein expression | NopT_USDA257_ | p5X-257T-F | agaacacgggggactctagatgctgcgtgatccgaacaat |
|  |  | p5X-257T-R | ccgctgttatcggtacctgtcaccttttgggtggtcaccg |
|  | NopT1_USDA110_ | p5X-110T1-F | agaacacgggggactctagaatgtatgatcgaatcggtgg |
|  |  | p5X-110T1-R | ccgctgttatcggtaccctgcatcctttgcgtcg |
|  | NopT2_USDA110_ | p5X-110T2-F | agaacacgggggactctagaatgtataatcgagtcgatggc |
|  |  | p5X-110T2-R | ccgctgttatcggtaccccgatgaggttccgc |
| Protein Expression in *E.coli* | NopT or NopT^C93S^ | 28a NopT-F | tgggtcgcggatccgaattcatgcacagtcccatcag |
|  |  | 28a FLAG-R | tcgagtgcggccgcaagcttcttgtcatcgtcatccttgtag |
|  |  | 28a Strep-R | tcgagtgcggccgcaagctttttttcaaattgaggatgagaccatcc |
|  | AvrPphB | ET28a-pphb-F | tgggtcgcggatccgaattcaaaataggtacgcaggccac |
|  | NopT_USDA257_ | ET28a-257T-F | tgggtcgcggatccgaattcctgcgtgatccgaacaa |
|  | NopT1_USDA110_ | ET28a-110T1-F | tgggtcgcggatccgaattctatgatcgaatcggtggctc |
|  | NopT2_USDA110_ | ET28a-110T2-F | tgggtcgcggatccgaattctataatcgagtcgatggcgaatac |
|  | NFR1^CD^ | duet NFR1-F | accacagccaggatccgagataccagaagaaggaagaagagaaagc |
|  |  | duet Sac1nost-R | ggcgcgccgagctcgcgatcggggaaattcgagct |
|  | NFR5^CD^ | ACSH-NFR5-F | agaagcgcggatccgaattctatgtatactgccgcagaaagaag |
|  |  | ACSH-NFR5-R | cgtatgggtagcttccaagcttacgtgcagtaatggaagtc |
|  | NFR5^KD^ | SUMO 5KD-F | ccgcgaacagattggtaaggttggggaatcagtgtac |
|  |  | SUMO HA-R | gctcgaattcggatcctttcacgcatagtcaggaacatcg |
|  | NFR5^JM^ | SUMO 5JM-F | ccgcgaacagattggttatgtatactgccgcagaaagaag |
|  |  | SUMO 5JM-R | gctcgaattcggatcctttcacgcatagtcaggaacatcg |
|  | sumo-NFR5^KD^-HA | duet sumo-F | catcaccacagccaggatgggtccctgcaggac |
|  |  | duet HA-R | cattatgcggccgcaagcttttacgcatagtcaggaacatcg |
|  | sumo-NFR5^JM^-GFP | duet sumo-F | catcaccacagccaggatgggtccctgcaggac |
|  |  | duet GFP-R | atgcggccgcaagcttttacttgtacaactcatccatacc |
|  | AtLYK5 | ACSH-LYK5-F | gcgcggatccgaattctacaaacgaaggtctaagaag |
|  |  | ACSH-LYK5-R | ggtagcttccaagcttgttgccaagagagccg |
|  | LjLYS11 | ACSH-LYS11-F | agaagcgcggatccgaattcacttgtctgaggaagagaaag |
|  |  | ACSH-LYS11-R | cgtatgggtagcttccaagcttacgagctgctatcaaagtt |
|  | LYK5^JM^ NFR5^KD^ | LYK5-NFR5-F | cgaaaacagaaaggttggggaatcagtg |
|  |  | NFR5 LYK5-R | ccccaacctttctgttttcgtcgctgaaatt |
|  | LYS11^JM^ NFR5KD | LYS11 NFR5-F | gagcagtgtaaggttggggaatcag |
|  |  | NFR5 LYS11R | ccccaaccttacactgctcattgagg |
|  | NFP^JM^ | ACSUG NFPJ-F | gtggaggcgctagaggatcctattgtctcaaaatgaagagattgaatagaag |
|  |  | ACSUG NFPJ-R | cttctcccttagagagctcacaattgtcactcagattcattg |
|  | NFR5^JM^-NFP^KD^ | 5JM-PKD-F | gatgagtgcaagattggtgaatcagtttacaaag |
|  |  | 5JM-PKD-R | accaatcttgcactcatcgct |
|  | NFP^JM^-NFR5^KD^ | PJM-5KD-F | gacaattgtaaggttggggaatcagtg |
|  |  | PJM-5KD-R | ccccaaccttacaattgtcactcagattcattg |
|  | NFR5^268-445^-NFP^458-595^ | 51/2KD-P1/2KD-F | ccatggccagaacttcaaccaactcaat |
|  |  | 51/2KD-P1/2KD-R | gaagttctggccatggcgaa |
|  | NFP^270-457^-NFR^5456-595^ | P1/2KD-51/2KD-F | ggatggctagaacttcgaccaac |
|  |  | P1/2KD-51/2KD-R | cgaagttctagccatcccgaa |
| site mutantions construction | NopT | T93S-F | gaatctccgtcggcttgactgc |
|  |  | T93S-F | gacggagattccatctgcgttc |
|  |  | 97D-F | ggcttggatgcggagtggctgcg |
|  |  | 97D-R | ctccgcatccaagccgacgcag |
|  |  | 97A-F | ggcttggctgcggagtggctgc |
|  |  | 97A-R | ctccgcagccaagccgacg |
|  |  | 109D-F | catccggatatccgaatggaggccctag |
|  |  | 109D-R | ttcggatatccggatgactgttgaggttacgc |
|  |  | 109A-F | catccggcaatccgaatggag |
|  |  | 109A-R | ttcggattgccggatgactgttg |
|  |  | 119D-R | cccggagatcaaaggcacgcctcagc |
|  |  | 119A-F | gcctttgatctccgggtactagggcc |
|  |  | 119A-F | cccggagcgcaaaggcac |
|  |  | 119A-R | ctttgcgctccgggtactagg |
|  |  | 156D-F | gcaaaacgatatgttgcagaaagcaggc |
|  |  | 156D-R | gcaacatatcgttttgcgccgcgaagtcggcc |
|  |  | 156A-F | gcaaaacgctatgttgcagaaagcag |
|  |  | 156A-R | ctgcaacatagcgttttgcgccgcgaagtc |
|  |  | H205A-F | ggcggcaacaccgttgcgacc |
|  |  | H205A-R | caacggtgttgccgccgccctcagc |
|  |  | D220A-F | ctcttcgctcctaatttcggcgaatttac |
|  |  | D220A-R | gccgaaattaggagcgaagagcgtggtgtttc |
|  |  | 245D-F | cgctacgacaatccaaaccggcag |
|  |  | 245D-R | ttggattgtcgtagcgattggctaggct |
|  |  | 257 mutants | changing the sequence of r primer at noptt257when cloned into vectors |
|  | NFR1 | K351E-F | gcaattgagaagatggatgtacaagcatc |
|  |  | K351E-R | catccatcttctcaattgctgttttcttgcc |
|  | NFR5 | 269-271A-F | ctatgcagctgcacgcagaaagaaggctctg |
|  |  | 269-271A-R | gcgtgcagctgcatagaattcggatccgcgc |
|  |  | 272-274A-F | ctgcgctgcagctaaggctctgaataggactgc |
|  |  | 272-274A-R | cttagctgcagcgcagtatacatagaattcggatcc |
|  |  | 275-277A-F | gaaaggcagctgcaaataggactgcttcatcagc |
|  |  | 275-277A-R | ctatttgcagctgcctttctgcggcagtatacatag |
|  |  | 278-280A-F | ctggctgcagctgcttcatcagctgagactg |
|  |  | 278-280A-R | agcagctgcagccagagccttctttctgcg |
|  |  | 282-283A-F | gcagcagctgagactgctgataaactac |
|  |  | 282-283A-R | gcagtctcagctgctgcagcagtcctattcagag |
|  |  | 285-286A-F | ggggctgctgataaactactttctggag |
|  |  | 285-286A-R | agtagtttatcagcagccccagctgatgaagcagtc |
|  |  | 288-290A-F | gctgcagcactttctggagtttcaggctatgtaag |
|  |  | 288-290A-R | aaactccagaaagtgctgcagcagcagtctcagctgatgaag |
|  |  | 291-294A-F | gagactgctgataaactagctgctgcagtttcaggctatgtaagcaagcc |
|  |  | 291-294A-R | atagcctgaaactgcagcagctagtttatcagcagtctcagctg |
|  |  | 295-298A-F | gagttgcagctgctgtaagcaagccaaacgtgtatg |
|  |  | 295-298A-R | cttacagcagctgcaactccagaaagtagtttatcagcag |
|  |  | 298-300A-F | ctatgcagctgcaccaaacgtgtatgaaatcgacg |
|  |  | 298-300A-R | gtttggtgcagctgcatagcctgaaactccagaaagtag |
|  |  | 301-303A-F | gcaaggcagctgcttatgaaatcgacgagataatggaagc |
|  |  | 301-303A-R | cataagcagctgccttgcttacatagcctgaaactcc |
|  |  | 304-306A-F | gtggctgcagctgacgagataatggaagctacgaag |
|  |  | 304-306A-R | cgtcagctgcagccacgtttggcttgcttacatagc |
|  |  | 307-309A-F | gaaatcgctgcagcaatggaagctacgaaggatttcag |
|  |  | 307-309A-R | ccattgctgcagcgatttcatacacgtttggcttgc |
|  |  | 310-312A-F | cgagatagcagctgctacgaaggatttcagcg |
|  |  | 310-312A-R | gtagcagctgctatctcgtcgatttcatacacgtttg |
|  |  | 313-315A-F | gctgcagctgcattcagcgatgagtgcaagg |
|  |  | 313-315A-R | ctgaatgcagctgcagcttccattatctcgtcgatttc |
|  |  | 316-318A-F | ggatgcagctgctgagtgcaaggttggggaatc |
|  |  | 316-318A-R | ctcagcagctgcatccttcgtagcttccattatctc |
|  |  | 319-321A-F | cgatgctgcagctgttggggaatcagtgtacaagg |
|  |  | 319-321A-R | caacagctgcagcatcgctgaaatccttcgtagc |
|  |  | 322-324A-F | gcaaggcagctgcatcagtgtacaaggccaacatag |
|  |  | 322-324A-R | gatgcagctgccttgcactcatcgctgaaatc |
|  |  | 325-327A-F | gaagcagctgcaaaggccaacatagaaggtcg |
|  |  | 325-327A-R | gcctttgcagctgcttccccaaccttgcactc |
|  |  | 268-277DE-F | agaagcgcggatccgaattcaataggactgct |
|  |  | 268-277DE-R | ctgatgaagcagtcctattgaattcggatccgcgcttc |
|  |  | 278-287DE-F | gcagaaagaaggctctggataaactactttctggagtttcagg |
|  |  | 278-287DE-R | aaactccagaaagtagtttatccagagccttctttctgcg |
|  |  | 298-307DE-F | caggctatgagataatggaagctacgaaggatttc |
|  |  | 298-307DE-R | cttccattatctcatagcctgaaactccagaaagtag |
|  |  | 308-317DE-F | gaaatcgacgatgagtgcaaggttgggg |
|  |  | 308-317DE-R | ttgcactcatcgtcgatttcatacacgtttggc |
|  |  | 318-327DE-F | gatttcagcaaggccaacatagaaggtcg |
|  |  | 318-327DE-R | gttggccttgctgaaatccttcgtagcttcc |
|  |  | 328-337DE-F | tcagtgtacgtaaagaaaatcaaggaaggtggtgcc |
|  |  | 328-337DE-R | attttctttacgtacactgattccccaaccttg |
|  |  | 288-294A-F | gcagctgccgcagcggctgcctcaggctatgtaagcaagcc |
|  |  | 288-294A-R | gcagccgctgcggcagctgcagcagtctcagctgatgaagc |
|  |  | 288-298PBS1-F | ggagacaaatctcatgtctcctcaggctatgtaagcaagcc |
|  |  | 288-298PBS1-R | ggagacatgagatttgtctccagcagtctcagctgatgaagc |
|  |  | N5-283S-R | tgaagcagtcctattcagagcc |
|  |  | 283Y-F | tctgaataggactgcttcatacgctgagactgc |
|  |  | N5-294G | agaaagtagtttatcagcagtctcag |
|  |  | 294Q-F | ctgctgataaactactttctcaggtttcaggctatgtaag |
|  |  | N5-303Y-R | cacgtttggcttgcttacatag |
|  |  | 304S-F | gcaagccaaacgtgtccgaaatcgacgag |
|  |  | N5-310AT-R | ttccattatctcgtcgatttcatacac |
|  |  | 311-2 IY-F | gaaatcgacgagataatggaaatctacaaggatttcagcgatgagtg |
| Complemental | NopT | HC-proT-F | gaattcgagctcggtacctcatggccttccttggagg |
|  |  | T-3-F | cacagtcccatcagtggttc |
|  |  | T-3-R | gaaccactgatgggactgtg |
|  |  | HC-T-R | cgcgggatcgagatctcgagtcatgtcatcttttgggtggtc |
|  | NFR5 | cherry-proNFR5-F | ccaagctgggctgcagggacatgagattgaagctcc |
|  |  | proNFR5-NFR5-F | ccccacttcacaaacatggctgtcttctttcttacctc |
|  |  | proNFR5-NFR5-R | gccatgtttgtgaagtgggg |
|  |  | NFR5-JM-KD-F | gatgagtgcaaggttgggg |
|  |  | NFR5-JM-KD-R | ccccaaccttgcactcatc |
|  |  | cherry kpn1 Sac1-R | ggcgcgcctaggtaccggggaaattcgagc |
